# DNA Methylation Ageing Atlas Across 17 Human Tissues

**DOI:** 10.1101/2025.07.21.665830

**Authors:** Macsue Jacques, Kirsten Seale, Sarah Voisin, Anna Lysenko, Robin Grolaux, Bernadette Jones-Freeman, Severine Lamon, Itamar Levinger, Carlie Bauer, Adam P. Sharples, Aino Heikkinen, Elina Sillanpaa, Miina Ollikainen, Cassandra Smith, James R. Broatch, Navabeh Zarekookandeh, Linn Gillberg, Ida Blom, Jesse Poganik, Mahdi Moqri, Vadim N. Gladyshev, Matthew Taper, Cassandra Malecki, Sean Lal, Nathalie Saurat, Steve Horvath, Andrew Teschendorff, Nir Eynon

**Affiliations:** Australian Regenerative Medicine Institute, Monash University, Clayton, Victoria, Australia; TruDiagnostics, Lexington, KY, USA; Institute for Health and Sport, Victoria University, Footscray, VIC, Australia; Novo Nordisk, Copenhagen, Denamark; Deakin University, Melbourne, Australia; Australian Institute for Musculoskeletal Science (AIMSS), Victoria University and Western Health, St Albans, Australia; Institute for Physical Performance, Norwegian School of Sport Sciences, Olso, Norway; Minerva Foundation Institute for Medical Research, Helsinki, Finland; Institute for Molecular Medicine Finland (FIMM), HiLIFE, University of Helsinki, Helsinki, Finland; Faculty of Sport and Health Sciences, University of Jyväskylä, Jyväskylä, Finland; Wellbeing Services County of Central Finland, Jyväskylä, Finland; Nutrition & Health Innovation Research Institute, School of Medical and Health Sciences, Edith Cowan University, Perth, Western Australia, Australia; Medical School, The University of Western Australia, Perth, Western Australia, Australia; Department of Biomedical Sciences, University of Copenhagen, Copenhagen, Denmark; Department of Biomedical Sciences, Faculty of Health and Medical Sciences, University of Copenhagen, Copenhagen, Denmark; Division of Genetics, Department of Medicine, Brigham and Women’s Hospital, Harvard Medical School, Boston, MA, USA; Faculty of Medicine and Health, The University of Sydney & Royal Prince Alfred Hospital; Altos Labs, Cambridge, United Kingdom; Shanghai Institute of Nutrition and Health, Chinese Academy of Sciences, University of Chinese Academy of Sciences, Shanghai, China

## Abstract

Aging involves widespread epigenetic remodeling across tissues, yet the nature and consistency of these changes remain unclear. We conducted a meta-analysis of more than 15,000 human methylomes spanning 17 tissues, identifying both conserved and tissue-specific aging signatures. We examined linear changes via differentially methylated positions, variability shifts via variably methylated positions, and Shannon-entropy to capture methylation disorder. Network analysis revealed fragile co-methylation modules largely resistant to beneficial perturbation. Key disruptors, including PCDHGA1, MEST, HDAC4, and HOX genes, exacerbated aging signals across tissues. Notably, a resilient module enriched for NAD□ salvage metabolism supports therapeutic targeting of NAD□ in aging. PCDHGA1 emerged as a conserved cross-tissue driver, suggesting protocadherin-mediated adhesion plays a broader role in maintaining structural and signaling stability in multiple organ systems. Our open-access atlas provides a foundational resource for dissecting the molecular architecture of human aging and identifying testable targets for intervention, biomarkers, and translational epigenetic therapies.

## Introduction

Aging is a universal yet profoundly individualized process. While chronological age progresses at a steady pace, its biological impact differs significantly among and within individuals, leading to varying risks of disease, functional decline, and mortality^1,2^. These differences are thought to stem from cumulative molecular changes, with epigenetic modifications—particularly DNA methylation, which involves the addition of methyl groups to DNA bases—as among the most reliable and consistent indicators of biological aging trajectories^1–3^.

Changes in DNA methylation associated with aging can typically be categorized into two main groups. The first group, differentially methylated positions (DMPs), consists of specific sites on the DNA that exhibit consistent changes in methylation levels across most individuals as they age, either increasing or decreasing in methylation. These changes are predictable and have been widely studied as potential biomarkers of biological aging^1–3^. The second group, variably methylated positions (VMPs), are sites on the DNA that display greater variability in methylation levels among individuals, and this variability tends to increase with age. Unlike DMPs, VMPs are less characterised across tissues and tend to not follow a uniform pattern across the population but instead reflect individual differences in epigenetic aging processes (i.e., environmental influences, genetic factors etc)^4–7^. These features capture both the deterministic components, which are predictable and influenced by specific biological processes, and the stochastic components, which involve random variations and noise within the aging methylome. This dual perspective helps in understanding the complex mechanisms underlying DNA methylation changes as organisms age^5,8^. Importantly, DMPs and VMPs are not mutually exclusive; a single CpG site can exhibit both a consistent age-related shift in methylation (DMP) and increased inter-individual variability with age (VMP), reflecting both directional and stochastic changes. For example, two individuals may both show age-related hypomethylation at a given CpG (a shared DMP), yet differ substantially in how stable or variable that hypomethylation is expressed across cells or time, capturing additional variation flagged by VMP analysis^7^. Entropy-based metrics, particularly Shannon-entropy, offer a quantitative lens into the complexity and disorganization of DNA methylation patterns across the genome. While DMPs contribute to age-associated increases in entropy by capturing cumulative directional shifts in methylation with chronological time, VMPs introduce baseline variability that raises entropy even in early life. Thus, entropy reflects both the accumulation of consistent age-related epigenetic drift (via DMPs) and the stochastic inter-individual variability (via VMPs) that exists independently of age. Elevated entropy values signal greater methylation heterogeneity, which is often interpreted as increased epigenetic noise and reduced regulatory precision^5,8,9^.

Although significant advancements have been made in our understanding of epigenetic changes associated with aging, most research has primarily focused on blood samples, with few specific tissue investigations available, resulting in a limited body of work in the field^7,10–13^. As a result, fundamental questions remain unresolved: to what extent is the aging methylome shared across tissues, and where does it diverge? Are common methylation signatures reflective of systemic aging, or are they tissue-restricted phenomena? And can epigenetic entropy and variability provide novel insights beyond traditional differential methylation? Understanding these dimensions is critical, as each tissue exhibits unique functional roles, cell-type compositions, and regenerative capacities that may shape distinct epigenetic aging trajectories. Solely focusing on blood risks overlooking tissue-specific alterations that could be key drivers of age-related dysfunction. At the same time, identifying conserved epigenetic patterns across tissues opens the door to using blood as a minimally invasive proxy to infer biological age in less accessible organs such as brain, heart, or muscle. This balance between shared and tissue-specific signatures is essential for refining biomarkers of aging and translating them into clinically meaningful insights.

To address these gaps, we assembled a comprehensive atlas of DNA methylation across 17 human tissues throughout the adult lifespan. Building on our detailed blood studies with over 30,000 participants^7^, we merged numerous open-source and collaborator datasets and performed integrated analyses across different organs. This work outlines the landscape of age-related DMPs, VMPs, and Shannon-entropy dynamics across tissues. This approach allows for the identification of universal ageing markers and tissue-specific differences, deepening our understanding of the molecular mechanisms of ageing. Additionally, using advanced computational tools, we have identified key drivers of ageing in different tissues and uncovered unique ageing pathways and coordinated methylation patterns among tissues.

Our atlas and newly built website (https://eynon-lab.shinyapps.io/humanagingatlas) serves as a valuable resource, providing a useful framework for understanding the signs of biological aging in human tissues. Additionally, our website, which showcases our results, acts as an essential reference for identifying biomarkers, acquiring mechanistic insights, and developing translational strategies to tackle human aging.

## Results

### A Pan-Tissue Landscape of Age-Associated DNA Methylation

To systematically characterize the aging methylome beyond blood, we assembled a cross-sectional atlas of DNA methylation across 17 human tissues, totalling over 15,000 samples from 131 datasets (**Supplementary Table 1**). We thoroughly curated these datasets and ensured quality assurance (see details in methods); only samples that passed our aforementioned quality control were included in the subsequent analysis. Prior to investigating age-related changes in the methylation, we have looked whether certain tissues had higher mean levels of methylation than others. Overall, global mean methylation ranged from 38% (cervix) to 63% (retina). However, when broken up by regions the distribution was much more uniform (**Supplementary Table 2**). Age-associated changes were evaluated using three key metrics: differentially methylated positions (DMPs), variably methylated positions (VMPs), and methylation Shannon-entropy, enabling a multidimensional view of epigenetic aging (**Supplementary Figure 1**). DMPs were identified using multivariate linear regression models that estimate changes in the mean methylation level at individual CpG sites concerning chronological age, while controlling for biological and technical covariates (e.g., sex, BMI, batch). In contrast, VMPs were identified using a two-step heteroscedasticity testing framework (Breusch–Pagan test), which detects age-associated differences in the variance of methylation levels, typically reflecting increased inter-individual variability in older individuals. While DMPs capture systematic, directional changes, VMPs reflect stochastic, instability-like effects that emerge with aging. Shannon entropy was used as a third, orthogonal metric to quantify the overall epigenetic complexity or disorder of methylation patterns at the sample level. Entropy values were derived from β-values adjusted for non-age covariates and calculated genome-wide, as well as across age-associated and non-age-associated CpGs. Shannon-entropy is maximized when CpG sites are approximately 50% methylated—indicating maximal uncertainty—and minimized when methylation levels are near 0% or 100%. By modeling entropy as a function of age, this analysis captures how the global unpredictability or stochasticity of DNA methylation evolves over time. Together, these three layers of analysis—mean-level shifts (DMPs), variance-level shifts (VMPs), and entropy-level disorder—provide complementary insights into how the methylome is remodeled with age at both locus-specific and system-wide scales. All three approaches were applied at the individual level within each dataset and then meta-analyzed within tissues, allowing the detection of age-related epigenetic patterns that are either tissue-specific, or across-tissues.

### Multivariate linear analysis of age reflects distinct tissue-specific aging signatures

We used multivariate linear regression models within each tissue, adjusting for relevant covariates, to identify CpG sites significantly associated with chronological age, thereby capturing tissue-specific DNA methylation aging signatures. The number of DMPs (i.e. age-associated CpGs) varied markedly across tissues (**Figure 1**). Brain, liver, and lung exhibited the largest numbers (e.g., brain: >180,000 DMPs), while pancreas, retina, and prostate showed minimal or no significant age-associations at an FDR < 0.005. Some tissues, such as skin and the colon, exhibited strong signatures despite moderate sample sizes. All tissues, except skeletal muscle and lung, showed an increased number of DMPs that became hypermethylated with age.

**Figure 1.**
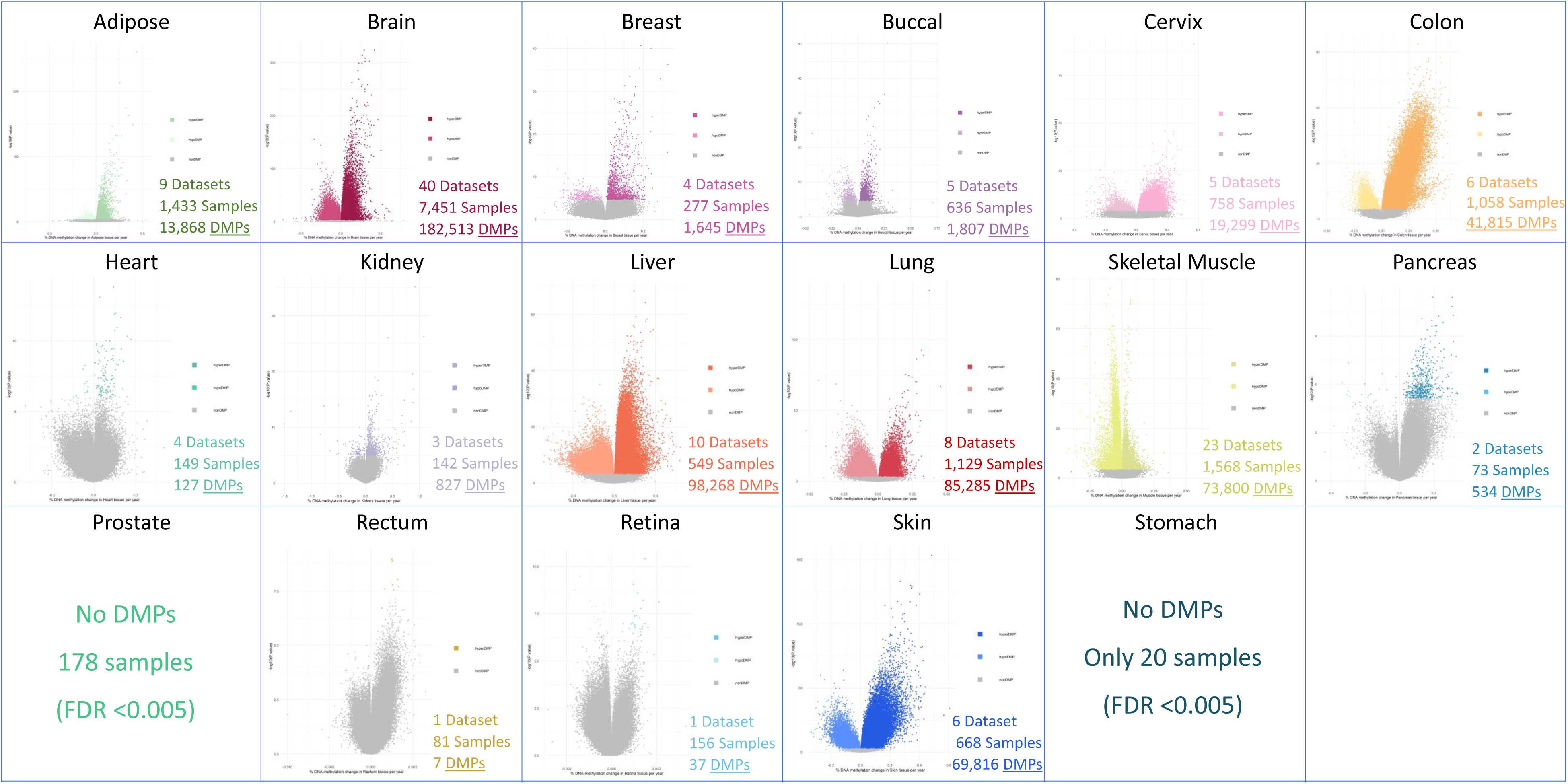
Differentially Methylated Positions (DMPs) with Age in 17 distinct human tissues. Each volcano plot depicts a unique tissue along with its methylation changes as one ages. Each dot corresponds to a distinct CpG, with colored dots indicating a significant association with age at an FDR <0.005, while gray dots signify CpGs that do not exhibit significant changes with age.

To further characterize the nature of age-associated methylation changes, we evaluated the distribution of methylation fractions (MF%) at DMPs between younger individuals (<30 years) and older individuals (>60 years) (**Figure 2**). In most tissues, age-DMPs were primarily found in regions with low methylation (<25%) in the younger group, with age-related hypermethylation being the predominant in the older group. For instance, in adipose tissue, 65% of DMPs in the young subjects were from low methylation regions, with approximately 70% of these reflecting positive (hypermethylated) differences between young and old individuals. This trend indicates a coordinated increase in methylation at previously unmethylated sites, potentially reflecting chromatin closure or the silencing of regulatory regions.

**Figure 2.**
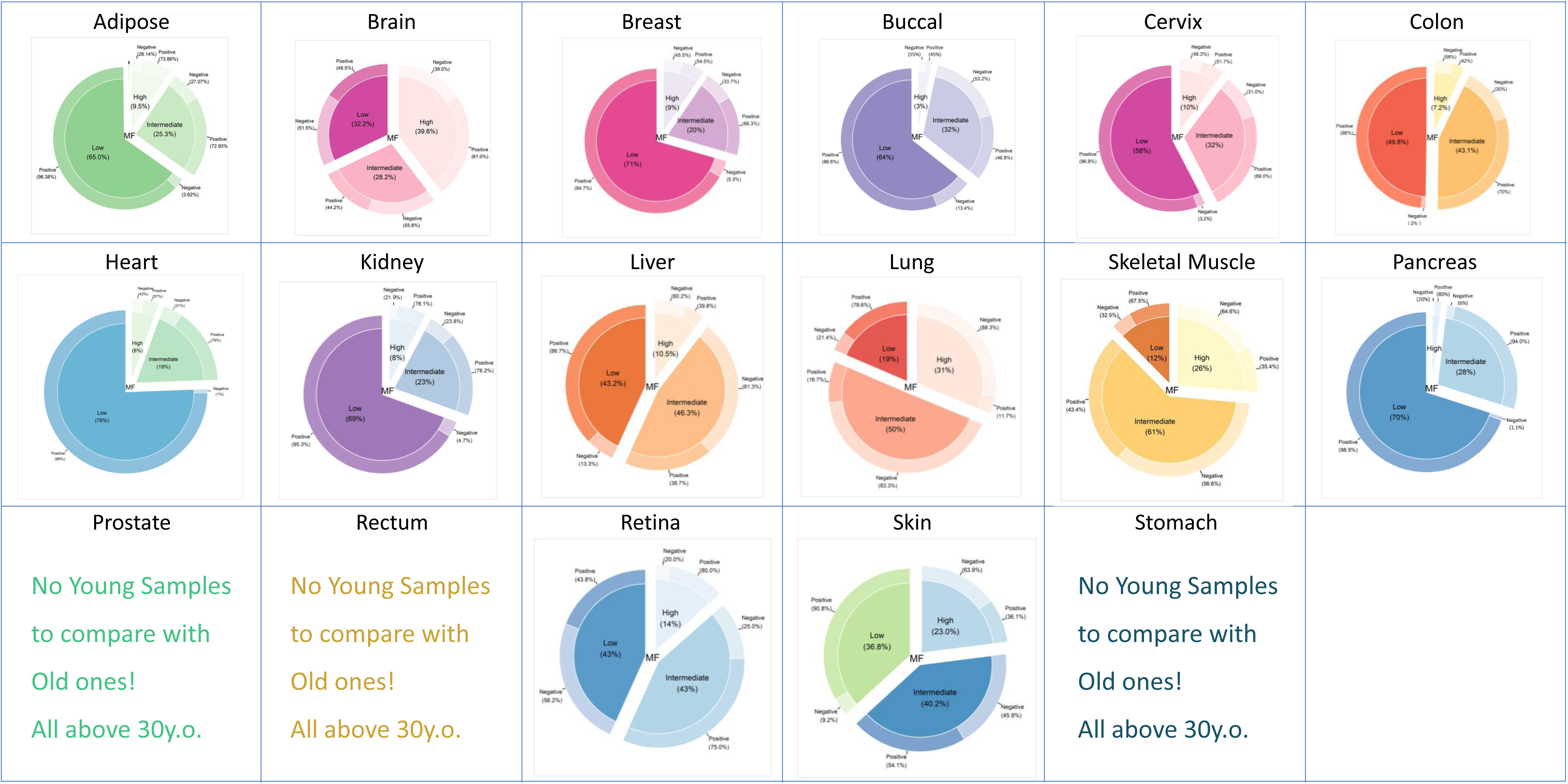
The fractions of Differentially Methylated Positions (DMPs) in young individuals compared to older individuals across 17 distinct human tissues. Each pie chart delineates a specific tissue alongside its corresponding methylation fractions for individuals aged less than 30 years and those aged over 60 years as they age. The inner circle signifies the percentage of DMPs classified as lowly methylated (<25%), intermediate methylated (25%-75%), or highly methylated (>75%) for individuals aged 30 years and younger. Conversely, the outer circle illustrates the methylation proportions within the population aged 60 years and older. For instance, in adipose tissue, a total of 13,868 DMPs associated with aging were identified. Of these DMPs, 65% were categorized as lowly methylated (beta values <0.25), 25.3% were classified as intermediate methylated (beta values between 0.25 and 0.75), and 9.5% were considered highly methylated (beta values >0.75) in younger individuals, as depicted in the inner circle of the adipose tissue pie chart. Analyzing the outer circle elucidates the alterations in DMP methylation status among individuals over 60 years of age. Specifically regarding adipose tissue, of the 65% lowly methylated DMPs identified in younger individuals, 96.38% exhibited elevated methylation levels, whereas 3.62% observed a decline. Among the 25.3% of intermediate methylated DMPs, 72.9% experienced an increase in methylation, while 27.07% demonstrated a decrease. Out of the 9.5% of highly methylated sites, 73.86% underwent an increase in methylation, and 26.14% experienced a decrease.

Other tissues exhibited a more balanced distribution among low, intermediate (25–75%), and high methylation (>75%) states. For example, tissues such as the colon, retina, liver, and skin showed a proportional distribution of DMPs (MF%) in intermediate methylation states (≥40%) and low methylation states (≥38%). With the exception of the retina, DMPs in low methylation states for these tissues generally showed hypermethylation states in aged individuals. Skeletal muscle and lung tissues exhibited 61% and 50%, respectively, of DMPs in the intermediate methylation state among young individuals. The majority of these sites showed hypomethylation in older individuals (83.3% in lung and 56.6% in skeletal muscle). Additionally, high methylation states in young individuals (31% in lung and 26% in skeletal muscle) were more likely to be hypomethylated in older individuals-specifically, 88.3% of lung DMPS and 64.6% of skeletal muscle DMPS in individuals aged 60 and above.

Brain tissue was identified as biologically distinct from all other tissues, showing a well-balanced distribution of DMPs across low, intermediate, and high methylation levels, all ranging from 28 to 40%. While there were age-related differences, they remained balanced, with hypermethylation and hypomethylation rates fluctuating between 40% and 60% in each fraction. Specific tissues, such as the prostate, rectum, and stomach, had insufficient samples from individuals under the age of 30 to be included in this analysis.

To assess the robustness of age-associated DMP detection across datasets, we performed a sensitivity analysis examining the relationship between the number of significant DMPs identified and key age-related characteristics of each tissue (**Supplementary Figure 2B**). Specifically, we evaluated whether the number of DMPs correlated with the mean age, median age, or age range of samples within each dataset. Among these, only age range showed a significant association with the number of DMPs, indicating that broader age distributions enhance the power to detect age-related methylation changes. This analysis supports the consistency and generalizability of our meta-analytic findings across diverse population structures.

### Variably Methylated Positions (VMPs) Are Scarce and Highly Tissue-Specific

Compared to DMPs, VMPs were much less frequent across tissues. Only a few tissues, such as brain, buccal, cervix, lung, and skin, had more than a handful of age-VMPs, whereas many other tissues showed few or no VMPs at the same FDR threshold (**Figure 3**). In general, the overlap of VMPs across tissues was limited, with fewer than 100 VMPs shared between any pair of tissues (**Supplementary Figure 3**). To determine whether the observed overlap of age-associated VMPs across tissues exceeded what would be expected by chance, we performed a permutation-based analysis. For each tissue comparison, we randomly selected sets of CpGs matched in number and genomic distribution to the observed VMPs and repeated this process 1,000 times to generate a null distribution of overlaps. The actual observed overlaps were then compared to this distribution to compute empirical p-values. This analysis revealed that half of the overlaps between tissue pairs occurred by chance, except for the pairs involving brain and buccal, buccal and cervix, buccal and lung, brain and skin, and buccal and skin (**Supplementary Table 3A).** To verify if the VMPs were detecting a biological signal rather than technical noise, we performed a correlation analysis between the number of VMPs and variables such as mean age, age range, and median range. VMP detection showed no significant correlation with age range, mean age, median age, or sample size (R ≈ 0.05, p > 0.8). This supports their tissue specificity and suggests that they are probably not artifacts of multiple study designs (**Supplementary Figure 2A**). These analyses collectively confirm that our atlas captures robust biological signals rather than technical noise.

**Figure 3.**
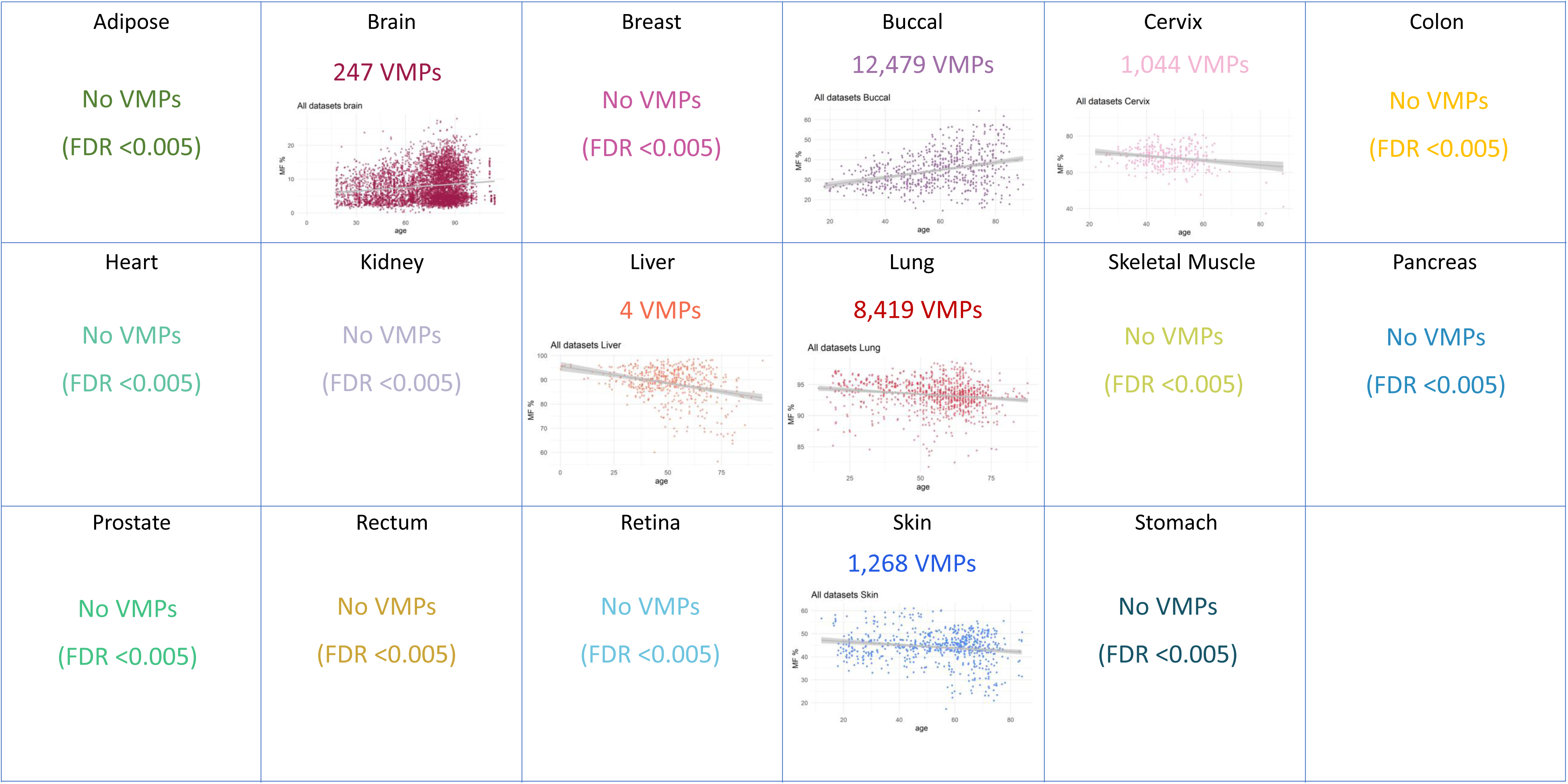
Age-related changes in variable methylated positions (VMPs) across 17 different human tissues. The scatter plot illustrates significant VMP changes specific to each tissue as individuals age. Each dot represents a unique individual.

Among the overlaps not attributed to chance, most overlapping VMPs showed similar trends, such as increases or decreases in variability (**Supplementary Figure 3**). However, in the case of Brain versus Buccal VMPs, only 15% of the 13 overlapping VMPs displayed similar directionality. We then conducted pathway enrichment analysis for both tissue-specific VMPs and overlapping VMPs. Most individual tissues revealed no enriched pathways for VMPs, except for Buccal and Lung tissues. Buccal VMPs showed enrichment across 210 pathways (FDR <0.005), particularly in areas related to system development (**Supplementary Table 4**). In contrast, lung VMPs were only enriched for three pathways: external encapsulating structure, extracellular matrix, and collagen-containing extracellular matrix. Based on this analysis, it is plausible to assume that VMPs occur randomly for most tissues with age, rather than in specific loci that are functionally linked.

### Shannon-Entropy and Epigenetic Disorder Reveal Tissue-Specific Aging Patterns

Entropy measures disorder in methylation and signal unpredictability. Entropy captures the overall disorder and unpredictability of DNA methylation patterns across the genome, offering a single quantitative measure that integrates all age-related differential changes— including those from low-variability DMPs. While variably methylated positions (VMPs) shift entropy toward higher or lower states, many of these CpGs are already hemi-methylated in youth, meaning entropy dynamics are not strictly age-dependent at these sites. This analysis therefore provides a complementary layer of insight beyond DMP or VMP-based approaches, revealing tissue-specific trajectories of epigenetic disorganisation and contributing to a broader understanding of methylation instability during ageing. (**Figure 4**). We observed that entropy increased preferentially at DMPs in several tissues such as adipose, breast, and kidney, suggesting a progressive loss of epigenetic fidelity at ageing-associated sites in multiple tissues. This pattern aligns with models proposing that epigenetic drift accompanies chronological ageing, particularly in metabolically active and/or proliferative tissues^5,7,14^.

**Figure 4.**
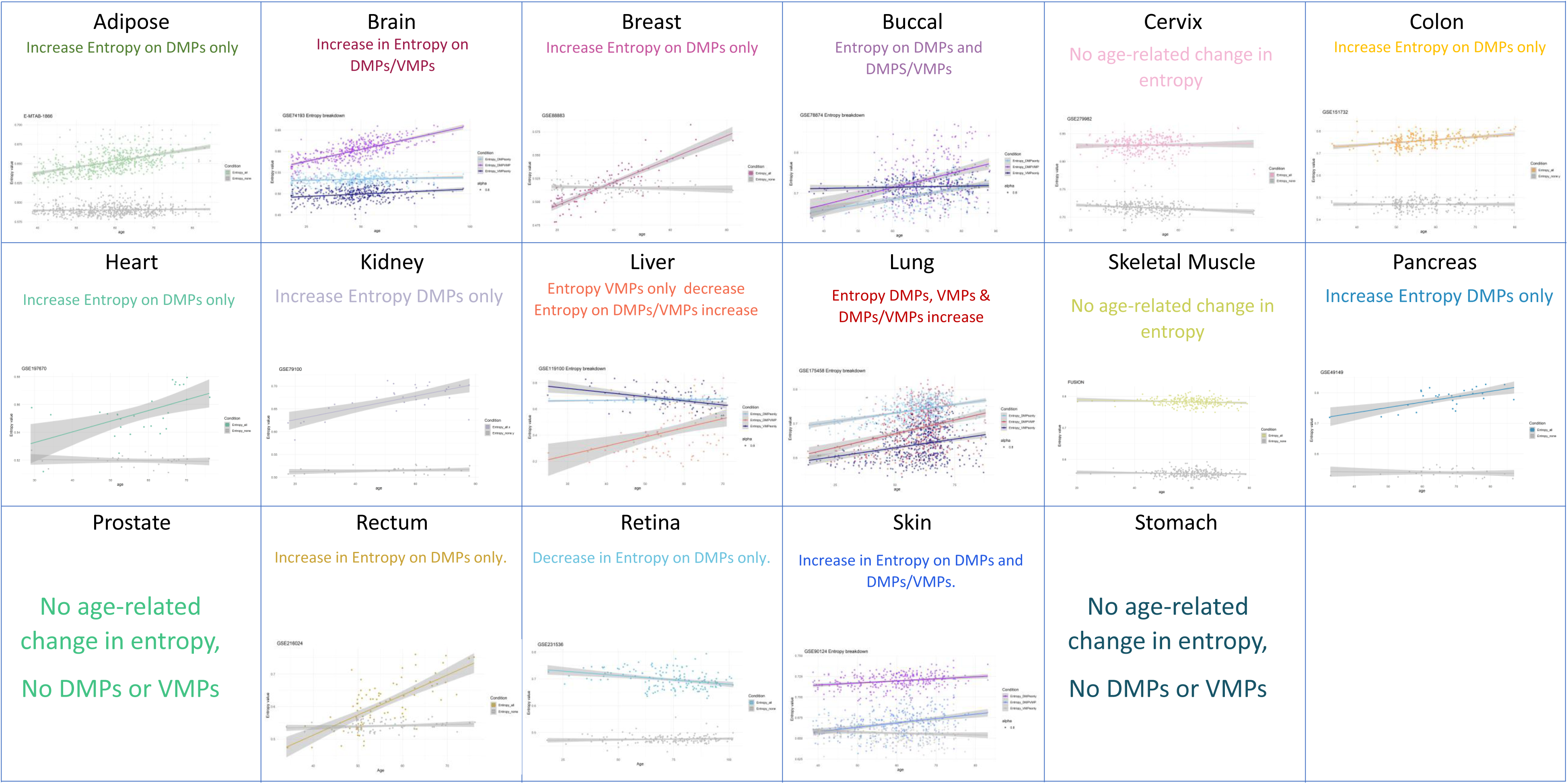
Entropy analysis plots were created for 17 different human tissues. Each plot represents specific tissues using the largest dataset available for each one. When variable methylated positions were observed in tissues, a division into DMPs only, VMPs only, and DMPs/VMPs was applied. Otherwise, a distinction between DMPs and non-DMPs with age served as the standard representation. Legends: Entropy_all: Entropy observed in age-DMPs. Entropy_none: Entropy observed in CpGs that are not DMPs for age. Entropy_DMPsonly: Entropy observed in age-DMPs only. Entropy_DMPVMP: Entropy observed in age-DMPs that are also age-VMPs. Entropy_VMPsonly: Entropy observed in age-VMPs only.

In contrast, other tissues, including the brain, buccal epithelium, liver, lung, and skin, exhibited entropy increases at positions that were both differentially and variably methylated (i.e., DMP–VMP overlaps). Such patterns indicate a more stochastic or plastic mode of epigenetic ageing in these tissues, where both directionality and variability of methylation are disrupted.

### WGCNA Highlights Tissue-Specific Hubs and Conserved Aging Modules

To gain clearer insight into aging-related methylation changes across various tissues and outline their co-regulatory framework, we conducted Weighted Gene Co-expression Network Analysis (WGCNA) on methylation data from selected tissues, including skeletal muscle, adipose tissue, blood (using a subset of samples as a proof of concept), and brain, using β-values to build tissue-specific co-methylation networks. Each tissue generated specific modules, with eigengene expression trajectories showing distinct aging signatures (**Supplementary Figure 4A/C/E/G**). Subsequently, we constructed a signed network and conducted hierarchical clustering on the topological overlap matrix to identify modules. The number and size of modules varied among tissues, reflecting biological complexity based on a data-driven approach (**Supplementary Figure 4B/D/F/H**). Module eigengenes, the first principal component of each module, were calculated and correlated with age to identify those with strong associations with aging. Significant positive and negative age-associated modules were included for further functional interpretation (**Supplementary Figure 5A/C/E/G**).

We enriched each module with Gene Ontology (GO) biological processes and categorized the enriched pathways according to module direction, allowing us to distinguish between age-related increases in methylation (positive modules) and decreases in methylation (negative modules). In adipose tissue, age-associated positive modules exhibited heightened methylation at loci related to forebrain development and cell adhesion, whereas modules with reduced methylation were enriched in synaptic signaling and NF-kappaB regulation, indicating a potential loss of regulatory control in immune and neuronal pathways (**Supplementary Figure 5B**). In blood, the CpGs in modules gaining methylation with age were related to embryonic development, cell-adhesion, apoptotic signaling, and immune pathways, while those losing methylation were connected to regulation and homeostasis (i.e. regulation of cell shape, regulation of body fluid levels) (**Supplementary Figure 5D**). The brain displayed increased methylation in modules enriched for GTPase activity, synaptic activity and organisation, cell communication and adhesion, among others, while decreased methylation impacted multiple morphogenesis-related processes and developmental pathways (**Supplementary Figure 5F**). In skeletal muscle, gains in methylation were observed in GTPase signaling pathways and some non-muscle developmental pathways. At the same time, losses were noted in pathways regulating muscle development and differentiation, and metabolic-related pathways (**Supplementary Figure 5H**). Beyond tissue-specific patterns, we also identified converging modules that revealed shared epigenetic aging programs across tissues. Notably, homophilic cell adhesion via plasma membrane adhesion molecules was enriched in adipose, blood and brain tissue (**Figure 5A/B/E**), implicating a common loss of cell–cell interaction integrity during aging. Meanwhile, small GTPase-mediated signal transduction was enriched in both brain and skeletal muscle (**Figure 5E/G**), pointing to conserved age-associated changes in intracellular communication and cytoskeletal regulation.

**Figure 5.**
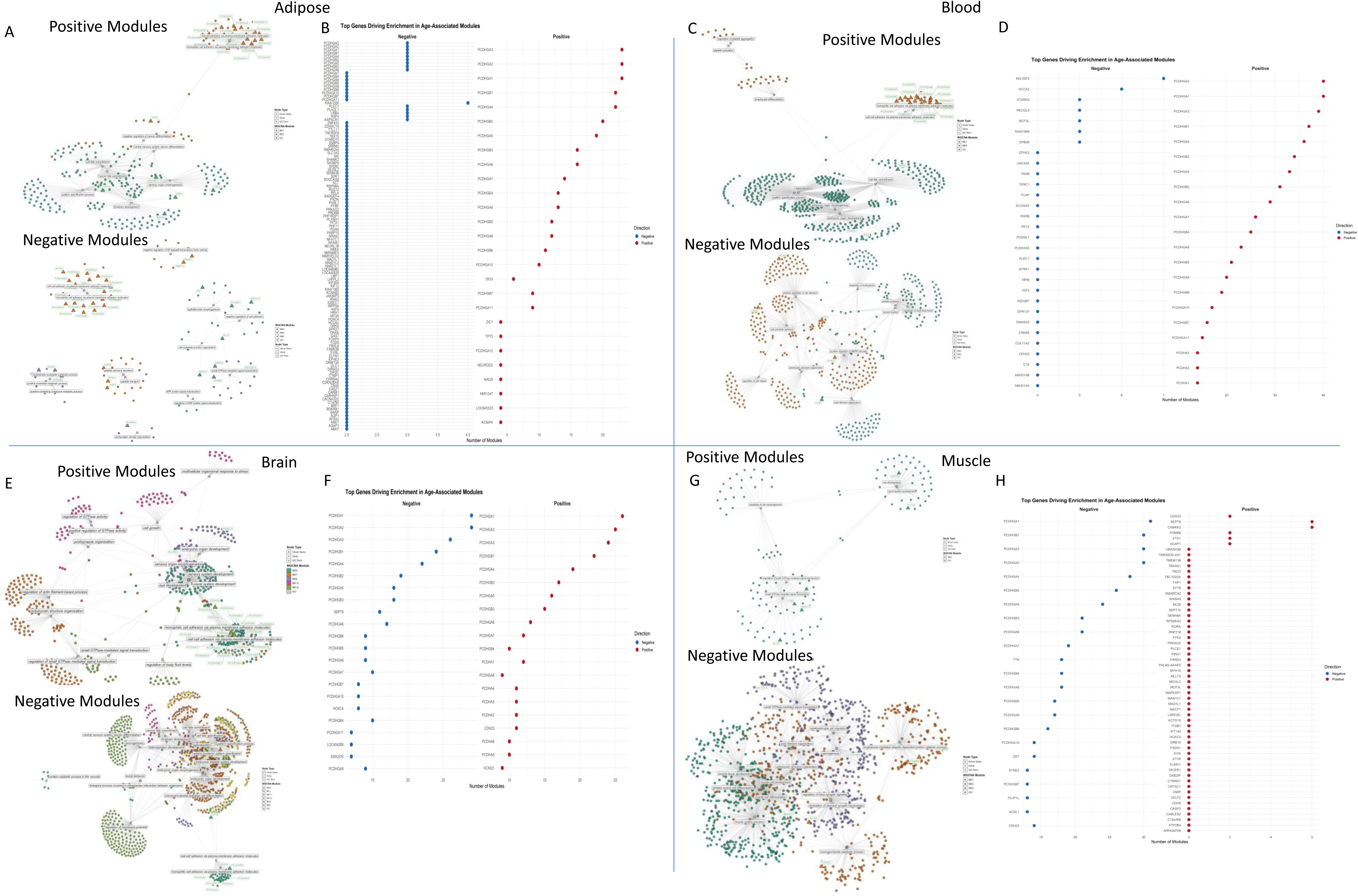
Weighted Gene Co-expression Network Analysis A/C/E/G displays the leading module pathways categorized by positive and negative modules. Each panel corresponds to a distinct tissue, with colors or dots indicating the module affiliation of these genes. Genes highlighted in green and represented as triangles in the networks are recognized as driver genes for each tissue module. The pathways associated with these gene groups are highlighted in gray. B/D/F/H show dot plots that illustrate the two genes responsible for enrichment in age-related modules. The y-axis indicates the gene names, whereas the x-axis reflects the number of pathways influenced by these genes within the modules.

WGCNA reveals module driver (hub) genes by examining intramodular connectivity. The CpGs correlating with genes demonstrating the highest association with the module eigengene (kME) are deemed essential for the module’s architecture and functionality (driver genes). Notably, all four tissues (adipose, brain, blood and skeletal muscle) identified Protocadherin Gamma (*PCDHG*) gene family members as primary module drivers (**Figure 5B/D/F/H**), a pivotal hub across various modules linked to cell adhesion and synaptic organization. Our results demonstrate that the consistent occurrence of *PCDHG* hub genes in diverse tissues indicates a conserved epigenetic responsiveness of protocadherin-mediated adhesion systems with the aging process. While traditionally linked to neural development and synaptic patterning, their persistent regulation through methylation in non-neural tissues, including blood and muscle, leads to the hypothesis regarding their broader function in preserving structural and signaling equilibrium as organisms age.

### Cross-Tissue Overlap Reveals Core and Divergent Epigenetic Aging Programs

To determine whether epigenetic aging signatures are consistent across tissues or predominantly tissue-specific, we analyzed the overlap of differentially methylated positions (DMPs) across all tissues in our atlas. We also included the blood DMPs identified in our previous blood atlas, publication^7^ . The intersection heatmap (**Figure 6A** – bottom diagonal half) depicts a complex scenario, showing that DMP overlap between tissues varies from 0% to 25% of the total DMPs, as determined by the tissue with the fewest DMPs. This suggests that most pairwise comparisons show a restricted number of common DMPs, emphasizing distinct aging pathways unique to each tissue, or possibly reflecting a reduced ability to detect DMPs, especially in tissues with limited samples. When evaluating the directionality of the intersected DMPs across tissues (**Figure 6A** – top diagonal half), a majority exhibit changes in the same direction, with direction concordance exceeding 59% for all tissue pairs except the brain and retina. The brain and retina demonstrated the highest level of discordant directionality, with only 28% of the 25 overlapping DMPs moving in the same direction. In our search for DMPs with the greatest overlaps among tissues, we identified cg16867657 as significant in 15 tissues (excluding the rectum). This specific CpG site is located on chromosome 6 at position 11044643-11044645, harbouring the well-known aging-related gene ELOVL2. A total of thirty-seven DMPs were identified across thirteen or more tissues. These DMPs included several genes that are well-established in the field of aging, as well as a few that are being linked to aging for the first time (**Table 1**).

**Figure 6.**
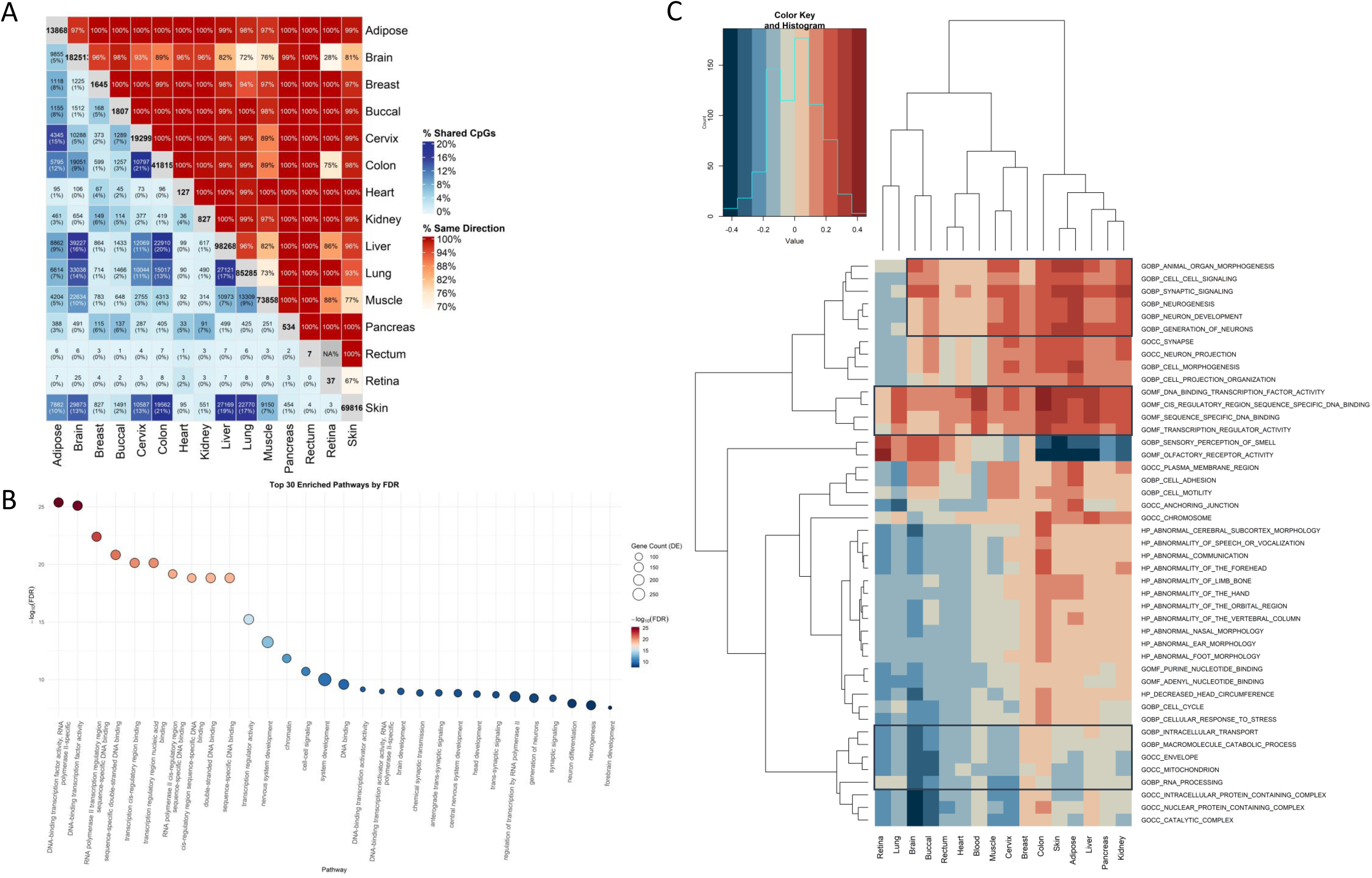
Cross-tissue overlap of age-associated DMPs reveals both conserved and tissue-specific aging signatures. A Bottom diagonal half of heatmap showing pairwise Jaccard indices of DMP overlap across human tissues profiled in the aging methylation atlas. Darker blue colors indicate greater overlap, with clustering revealing groups of tissues with shared age-associated DMPs. Most tissues show low overlap, reflecting divergent aging trajectories. Top diagonal half of heatmap shows effect-size correlation across tissues to determine directionality changes. Darker red colors indicate greater concordance in direction of changes of overlapped DMPs. B Enrichment pathways identified from GpGs found in nine or more tissues. The colors of the dots indicate the FDR strength, while the circle sizes reflect the number of genes associated with each pathway. C Heatmap generated from the *mitch* analysis. This heatmap depicts the 50 most enriched pathways revealed by the mitch analysis. Blue indicates hypomethylation of genes within these pathways, whereas red signifies hypermethylation of genes in the pathways. The enriched pathways are listed on the y-axis, while the x-axis shows the results for each tissue.

**Table 1.** Top pan-tissue DMPs present in 13 or more tissues.

To ensure that the observed overlaps were not merely due to random chance, we performed permutation tests as described in the VMPs section, by randomly rearranging the sample age labels within each tissue and recalculating DMP overlaps over 1,000 iterations. As shown in **Supplementary Table 3B**, the results indicated that the empirical overlaps between several pairs of tissues significantly surpassed the 95th percentile of the null distribution, providing compelling evidence against coincidental concordance. The exceptions were buccal vs. rectum, cervix vs. retina, liver vs. retina, rectum vs. retina, retina vs. blood and retina vs. skin. Notably, only one dataset was found in the analysis of the retina and rectum, which significantly limits the ability to identify age-related changes in these tissues, likely resulting in observed overlaps that do not meet the null distribution.

Finally, we conducted a sensitivity analysis to confirm that our DMPs’ identification was not influenced by technical artifacts such as age ranges, mean age, or median age. The updated scatterplots examining DMPs versus age (**Supplementary Figure 2B**) illustrate our tests for correlations between the number of DMPs and the age range, median age, and mean age across different tissues. Our analysis revealed that only the age range demonstrated a significant correlation with DMP count (R = 0.66, p = 0.0072), while mean and median age showed no significant correlation (R = 0.13 and 0.22, respectively). This suggests that a broader sampling across the lifespan enhances detection power without introducing systematic bias.

### Cross-Tissue Network Level Analysis through Module Enrichment Identifies Key Pathways and Genes as Drivers of Aging

We subsequently investigated the functional significance of genes that consistently showed age-related changes in methylation across various tissues. We established a rigorous consensus set of genes differentially methylated in nine or more tissues (>50% overlap across tissues) and conducted pathway enrichment analysis on this core group. Notably, a prominent signature surfaced: almost all enriched pathways were associated with transcriptional regulation, chromatin organization, and gene regulation (**Figure 6B**).

While compelling, this method has constraints due to statistical power; tissues with fewer samples or minor methylation changes may not be adequately represented (**Supplementary Table 1**). Additionally, it restricts interpretability as it does not enable comparison of directionality across tissues in each pathway. To tackle these issues, we employed *mitch*^15^, which combines the magnitude and direction of gene-level shifts across different tissues using t-statistics, rather than depending only on significance thresholds. This approach allowed underpowered tissues to meaningfully contribute to the overall analysis landscape. The integration based on *mitch* essentially validated the transcriptional regulation signature while broadening it to include additional pathways relevant to aging, such as mitochondrial function, cell-cycle pathways, and DNA damage response (**Supplementary Table 5**). A thorough examination of the top fifty pathways has unveiled three distinct clusters that illustrate a consistent shift in agreement across nearly all tissues (**Figure 6C**). For instance, pathways associated with transcription factor activity, as observed in the analysis above, DNA binding, and transcriptional regulation were identified as being enriched in hyper-DMPs across all tissues. Additionally, pathways related to cell-cell signalling, synaptic signalling, and neurogenesis were observed to be enriched in hyper-DMPs across all tissues, with the exception of the retina and lung. Conversely, pathways involved in intracellular processes, such as those pertaining to mitochondria and macromolecule metabolism, were found to be enriched in hypo-DMPs in all tissues, except for breast and colon.

To further investigate the signatures identified by *mitch*, we generated a gene–gene correlation matrix across tissues utilizing Spearman correlation. This method evaluated the consistency of each gene’s aging profile across various tissues. For instance, if two genes increase in methylation levels with age in most tissues, they are considered to have a strong positive correlation. Conversely, if the methylation of one gene increases in some tissues while in another decreases, the correlation is regarded as negative. Consequently, we were able to model “t-patterns,” indicating whether a gene is uniformly upregulated or downregulated with age across tissues, or exhibits fluctuations that are specific to certain tissues. We utilized hierarchical clustering and dynamic tree cutting on the gene–gene correlation matrix to extract biological structures from these patterns, grouping genes into modules based on co-aging behaviour. Unlike WGCNA modules, these modules arise from cross-tissue coherence in aging trajectories as indicated by t-statistics, rather than solely from co-methylation patterns in a single tissue.

Each resulting module captures a group of genes that age uniformly across different tissues. Some modules exhibited significantly high positive t-scores in nearly all tissues, indicating a widespread hypermethylation linked to aging (**Supplementary Table 4**). In contrast, other modules were distinctly tissue-specific, showing aging-related hypermethylation in one tissue while demonstrating hypomethylation in another, suggesting varying aging mechanisms. This analysis uncovered several important findings: for instance, Module 2 showed elevated average t-statistics in blood, indicating significant hypermethylation associated with aging, while also revealing negative t-statistics in the brain, suggesting hypomethylation. Only one module (Module 5) exhibited a universal pattern, indicating hypermethylation across all tissues (**Supplementary Table 4, Supplementary Figure 6**). Importantly, the driver score gene in this module belongs to the *PCDHG* gene family (*PCDHGA1*), reinforcing the idea that this gene family may play a key role in aging processes throughout all human tissues. In summary, our findings highlight that specific gene programs (module clusters) are regulated in a highly tissue-specific manner during aging, while others may indicate shared core aging mechanisms. This may also present tissue-specific or even contradictory regulations (e.g., beneficial in one tissue but harmful in another).

### In-Silico Robustness Analysis Reveal Functionally Fragile Genes Driving Module Architecture

To advance from correlation to testing functional effects, we established an in-silico validation framework. In this framework, we computationally adjusted methylation levels for each gene, simulating conditions like hypermethylation, and reassessed the structural integrity of aging-associated modules. This simulation-driven strategy identified a limited set of “global disruptor” genes that modified the arrangement of various modules across different tissues (**Supplementary Table 5**). The foremost disruptor genes encompassed the PCDHG gene family, MEST, HDAC4, and genes from the HOX family. When combined with additional key genes from each module, they instigated significant changes in modular connectivity, with some modules exhibiting alterations exceeding 50% when key genes were modified (**Supplementary Table 5**). Considering the enrichment pathways of each module, it is reasonable to assert that alterations in DNA methylation result in changes in function pertinent to these pathways.

To better understand the biological effects of these disruptions, we integrated module-level enrichment findings and classified gene modifications as potentially beneficial or detrimental. For instance, if a gene disrupts a module linked to aging pathways and disruption diminishes the module’s influence, it is deemed beneficial; conversely, if the disruption enhances the module’s effect, it is considered harmful. Only ten modules presented a beneficial disruption, while all others appear to have a harmful effect if disrupted. These findings suggest that most aging-related modules are vulnerable to modifications, with only a (**Supplementary Table 5**) small subset showing potential for beneficial modulation.

## Discussion

Our cross-tissue atlas, which includes over 15,000 methylomes derived from 17 human tissues, presents one of the most comprehensive pictures to date of the cross-tissue functional epigenetic alterations associated with human aging. Through a thorough analysis of three layers of methylome dynamics, DMPs, VMPs, and Shannon-entropy, we have shown that aging is associated with both tissue-specific and conserved cross-tissue effects on the methylome. The heterogeneity of DMPs across various tissues highlights the complex and organ-specific nature of the aging process^9,16,17^.

Tissues such as the brain, liver, lung, skeletal muscle, and skin exhibited a large number of DMPs associated with age, suggesting a reconfiguration of the methylome or vulnerability associated with advanced age^5,18,19^. In contrast, tissues such as the kidney, prostate, rectum, and stomach demonstrated minimal to no detectable changes, likely attributed to the limited sample size or an inherently more stable epigenetic landscape. This is evidenced by the fact that comparable studies in the domain have identified a significant number of DMPs associated with age, even within smaller datasets^18^. More data is required in these rare tissue types to confirm this hypothesis. Despite this tissue variability, a discernible pattern emerged: most tissues demonstrated age-associated hypermethylation, particularly in previously unmethylated regions in younger individuals^18–20^. This observation implies a transition toward epigenetic silencing, likely resulting from the closure of open chromatin regions, affecting enhancer activity or gene expression regulation^20,21^. Exceptions to this trend, were skeletal muscle and lung tissues, which demonstrated heightened hypomethylation, may signify tissue-specific aging mechanisms^22–24^, encompassing demethylation at regulatory or structural loci.

Building on these DMP patterns, we next sought to determine how these age-associated DMPs shift across the methylation spectrum. In multiple tissues, regions that exhibited low methylation levels (<25%) in young individuals showed a marked increase in methylation with age. Similarly, DMPs with intermediate methylation levels (25–75%) in youth often showed higher methylation in older individuals. This common observation of higher methylation in older individuals suggests that, in several tissues, a trend toward global hypermethylation is more pronounced than one toward stochasticity or increased epigenetic entropy—contrary to the often-assumed paradigm of random drift toward intermediate methylation states in aging^9,17,25^. In some instances, even DMPs that were already highly methylated (>75%) in young individuals displayed even higher methylation levels in older individuals, reinforcing the concept of progressive chromatin condensation and regional silencing as a potential aging mechanism ^3,26^. Certain tissues, like the colon, liver, and skin, showed a more even distribution of low and intermediate methylation levels among the youth. In these instances, age-related changes were more varied, indicating both hypermethylation and hypomethylation based on the initial methylation status. This trend may suggest tissue-specific regulation or differing sensitivities to age-related environmental factors. Given the known characteristics of these tissues, it is reasonable to hypothesize that such patterns are influenced by cell turnover rates, metabolic activity, or immune infiltration^27,28^. Further investigations into this area are recommended as the next steps.

The brain emerged as epigenetically distinctive, characterized by an equitable distribution of DMPs across various methylation states and a bimodal directionality of change. This phenomenon may reflect a more sophisticated or buffered methylation landscape, aligning with the cellular diversity of the brain and its relatively low mitotic activity^29^. This balanced pattern of methylation changes in the brain might also result from analyzing multiple brain regions together. Since each region likely has its own unique epigenetic signature, the overall pattern may reflect an average of different cell-type-specific changes, rather than changes unique to the brain as a whole^14^. Importantly, these findings add a layer of directional nuance to the epigenetic aging narrative, demonstrating that age-related methylation changes are not solely characterized by increased noise or drift, but also by systematic and tissue-dependent remodeling of the methylome, including progressive reinforcement of silenced states. This insight expands upon previous aging epigenome studies that primarily quantified methylation change as a binary gain or loss, and highlights the importance of baseline methylation state in shaping the trajectory of age-associated epigenetic remodeling. Future studies should investigate whether these directional shifts correspond with chromatin accessibility and histone modifications, and whether they confer functional consequences on regulatory landscapes relevant to tissue aging and disease vulnerability.

In contrast to DMPs, variably methylated positions (VMPs) were scarce and exhibit a high degree of tissue specificity, supporting the hypothesis that aging-related epigenetic drift transpires in a stochastic and localized manner^5,7,30,31^. Only a limited number of tissues demonstrated a significant number of VMPs. Furthermore, VMPs predominantly exhibit a lack of consistent pathway enrichment across the majority of tissues, with the exception of the buccal and lung tissues, indicating that these may represent arbitrary degradations of epigenetic regulation rather than orchestrated biological responses^32^. This finding challenges the prevailing assumption that increased variability in methylation is uniformly detrimental or functionally significant in the context of aging, thereby highlighting the necessity for a more nuanced framework that effectively differentiates noise from signal^30^. Entropy, frequently acknowledged as a hallmark of aging-associated molecular chaos, is tissue-dependent and closely correlated with metabolically active organs. Adipose, breast, and kidney tissues displayed the highest increases in entropy at canonical DMPs, potentially reflecting a progressive loss of regulatory control in energy-demanding systems^33,34^. Notably, the entropy profile of the brain was uniquely balanced across DMPs and VMPs, suggesting the existence of a distinct form of methylomic aging that may mitigate both directional and stochastic changes^14^.

Our WGCNA-based analysis uncovered modules of co-methylated CpGs with distinct aging trajectories and functional enrichments. Modules showing higher methylation in older individuals tended to be enriched in developmental and adhesion pathways, whereas those with decreased methylation were enriched in immune, metabolic, and neurogenic signaling. This duality mirrors known hallmarks of aging—loss of regenerative capacity and altered intercellular communication^1,2^. The consistent enrichment of GTPase signaling and cell adhesion pathways across multiple tissues suggests that aging recurrently impacts biological programs governing synaptic structure and cytoskeletal integrity. Recent research underscores that small GTPases—such as members of the Rho, Ras, and Rab families—play central roles in maintaining cytoskeletal dynamics, vesicle trafficking, and cell communication, functions that decline with age^35,36^. Notably, dysregulation of GTPase activity has been implicated in neurodegeneration, immunosenescence, and cellular senescence^36^. The recent increase in interest in creating pharmacological agents that influence GTPase activity as potential gerotherapeutics^37,38^ aligns with our findings, providing indirect evidence of the validity of our results.

A notable finding was the repeated identification of Protocadherin Gamma (PCDHG) family genes as key drivers in both WGCNA tissue-specific networks and the pan-tissue analysis. These genes are crucial for cell adhesion and synaptic organization^39^, suggesting a role in maintaining structural integrity during aging. Recent research links hypermethylation in the PCDHG family to reduced white matter in the brain^40^, a marker of accelerated cognitive decline. This connection is particularly intriguing since previous studies have shown that abdominal fat can predict decreases in cerebral white matter^25,41^. This gene family is also associated with various age-related diseases, including increased inflammation in bronchial epithelial cells^42^, reduced stroke volume and ventricular dysfunction^43^, pre-cancerous gastric lesions^44^, colorectal cancer^45^, hepatocellular carcinoma^46^, and age-related skeletal muscle weakness^47^. Identified as key drivers across all four tissues examined by WGCNA and consistently in our pan-tissue analysis, we propose that modifications to the DNA methylation patterns of these genes may adversely affect age-related pathways. Our in-silico studies indicate that manipulating this gene family impacts 100% of its related module, functionally influencing several aging pathways. In conclusion, the relationship between these genes and aging-related diseases across various tissues, corroborated by earlier studies^25,40–47^, combined with our pan-tissue analysis, which identifies these genes as central to aging, and our in-silico disruption results that highlight their role in epigenetic reprogramming associated with aging, suggests that the tissue-specific methylation patterns of this gene family could serve as biomarkers for aging and warrant further investigation as targets for anti-aging therapies.

While many DMPs are unique to specific tissues, several CpGs—such as in genes ELOVL2, KLF14, FHL2, TBR1, and TRIM59—identified as aging biomarkers in individual tissues, have also emerged as pan-tissue aging markers in this study. This not only highlights their importance as dependable biomarkers of aging but also broadens their impact as indicators for the entire body, confirming the robustness of our analysis findings. Permutation tests validated that these overlaps were unlikely to be due to chance, particularly in tissues with larger sample sizes. Interestingly, DMPs often showed conserved directionality across tissues, even when overlaps were minimal. This tension between global conservation and local specificity could underpin tissue-level manifestations of systemic aging, such as frailty or multi-organ failure^48^. Since many of our pan tissue genes are recognized as biomarkers of aging in individual tissues, we propose that genes which have been less studied in the aging context and demonstrate consistent directionality across tissues could also act as systemic biomarkers. In contrast, discordant CpGs between different tissues may indicate adaptive or compensatory responses.

Our mitch-based meta-analysis expanded the picture by integrating gene-level statistics across tissues. This revealed broad upregulation of pathways tied to transcriptional regulation and chromatin remodeling (e.g., DNA binding, histone modification), as well as age-related suppression of metabolic and mitochondrial processes. Notably, transcriptional repression via DNA hypermethylation emerged as a conserved hallmark, while intracellular metabolic decline via hypomethylation was evident in all tissues except breast and colon. This suggests an imbalance between nuclear and cytoplasmic aging programs, a theme increasingly recognized in aging biology^49^.

By constructing a gene–gene correlation matrix using tissue-level t-statistics, we identified groups of genes in which the magnitude of change showed either coordinated or discordant methylation shifts across tissues. This allowed the construction of cross-tissue modules that more faithfully represent multi-organ aging trajectories than WGCNA alone. Some modules showed strong coherence across tissues (e.g., Module 5), while others revealed tissue-specific divergences, supporting the idea of both universal and compartmentalized aging programs. To evaluate functionality and vulnerability, we simulated gene-level methylation changes and assessed their impacts on module structure. Genes like *PCDHGA1, MEST, HDAC4*, and members of the HOX family surfaced as global disruptors, indicating that their altered methylation significantly reshaped aging modules across different tissues. Most disruptions were harmful, as they exacerbated aging markers, although a few alleviated these effects. This analysis identified only 10 modules with beneficial disruptions, indicating that the aging methylome largely revolves around fragile, pro-degenerative networks with limited opportunities for therapeutic intervention. However, these few disruptable modules present key therapeutic prospects. For example, one of the leading modules identified (Module_160) exhibited remarkable resilience, with its top five beneficially modulated pathways including NAD□ biosynthesis through the nicotinamide riboside (NR) salvage pathway. This finding directly correlates with an expanding body of research highlighting the importance of NAD□ metabolism in healthy aging and longevity^50^. NAD□ levels decrease with age, leading to mitochondrial dysfunction, decreased sirtuin activity, and heightened DNA damage. Numerous studies indicate that restoring NAD□ levels, especially through precursors like NR or nicotinamide mononucleotide, can counteract aspects of age-related decline in various tissues, including the brain^51^, muscle^52^, and heart^53^, and can prolong healthspan in model organisms^51,54–56^. The identification of the NAD□ salvage pathway as one of the most significantly modulable nodes in our analysis emphasizes its crucial role in maintaining cellular balance and resilience during aging, providing epigenetic evidence to support its therapeutic targeting.

### Conclusions and Limitations

In conclusion, our study presents a comprehensive atlas of human epigenetic aging, integrating over 15,000 methylomes across 17 tissues and capturing multiple layers of methylation dynamics—differential methylation, variability, and entropy. This atlas reveals that aging is not solely characterized by stochastic drift but involves systematic, tissue-specific, and often directional methylome remodeling, with consistent enrichment of pathways tied to development, cytoskeletal integrity, and immune regulation. Importantly, our cross-tissue and in silico analyses identify both fragile modules that exacerbate aging signatures and rare, resilient modules—such as those enriched for NAD□ biosynthesis—that offer therapeutic promise. As such, this work provides a powerful, openly usable resource for the academic community to explore aging mechanisms (https://eynon-lab.shinyapps.io/humanagingatlas), prioritize biomarkers, and identify candidate pathways for intervention.

Nevertheless, certain limitations should guide interpretation. Unequal sample sizes across tissues may limit detection power in underrepresented organs (**Supplementary Table 1**), and furthermore, the use of bulk tissue masks cell-type-specific signals and compositional shifts. The cross-sectional nature of the data constrains our ability to capture temporal trajectories of epigenetic aging, and the absence of complementary omics layers, such as chromatin accessibility or gene expression, limits mechanistic insight. Although computational predictions offer testable hypotheses, experimental validation is essential to confirm causal roles for identified genes and modules. Despite these challenges, our atlas sets the stage for deeper, integrative studies of the aging methylome, and offers a scalable framework for understanding and potentially modulating human aging at the epigenetic level.

## Methods

### Study Design and Scope

This study represents a large-scale, cross-tissue epigenome-wide association meta-analysis (EWAS) of human aging, integrating over 15,000 methylomes from 17 different tissues. Utilizing advanced bioinformatics approaches, we systematically analyzed and interpreted extensive existing epigenomic datasets. Given the typically small effect sizes of age-related DNAm shifts, our strategy leveraged the statistical power of meta-analysis to overcome these limitations and enhance reproducibility. The methodological framework was especially suited for detecting subtle, yet consistent, methylation changes associated with chronological aging across diverse cohorts, including variations by sex and health status. Our approach expands on prior frameworks^7^ by not only identifying differentially methylated positions (DMPs), variably methylated positions (VMPs), and entropy dynamics, but also by incorporating advanced co-methylation network analyses (WGCNA), functional gene-level pathway meta-analysis (mitch), gene-gene pan-tissue correlation networks, as well as in-silico validations. These multilayered methods collectively offer a robust, systems-level perspective on epigenetic aging.

### Data Collection and Pre-processing

We assembled a large repository of DNAm profiles encompassing 15,995 human samples from 131 independent datasets, each assayed using Illumina methylation arrays (27K, 450K, EPIC). Data sources included 112 open-access studies from GEO, 5 from ArrayExpress, 2 from dbGaP, one from EGA, and 10 from independent collaborators. Datasets with fewer than 10 samples or low age dispersion (standard deviation <5) were excluded to preserve statistical robustness. Additionally, samples from individuals diagnosed with cancer were removed to avoid confounding due to aberrant methylation patterns. For datasets with available raw intensity data (IDATs), pre-processing and normalization were conducted in R using the *ChAMP* pipeline^57,58^. Quality control filters were applied to remove samples with >10% of probes failing detection (p□>□0.01), as well as probes with missing β-values, low bead count, non-CG content, cross-hybridization, or mapping to SNPs or sex chromosomes in mixed-sex datasets. Normalization of type I and II probes was done via champ.norm, and unwanted technical variation (e.g., slide, position effects) was adjusted within each dataset using *ComBat*^59^ when metadata was available. Sex mismatches between annotation and predicted sex were resolved using getSex from the *minfi* package.

### Differential Methylation Analysis (DMPs)

To identify age-associated differentially methylated positions (DMPs), linear models were fitted independently within each dataset using *limma*^60^. Models incorporated available covariates such as sex, BMI, and technical batch (**Supplementary Table 1**). Where applicable, repeated or related samples (e.g., twins) were modeled using random effects via duplicateCorrelation. To decide on covariate inclusion in the models, we referenced GBD data and incorporated all non-are-related covariates, such as asthma, anemia, depression, etc. Age-related diseases like Alzheimer’s, stroke, T2D, and CVD were excluded. Additionally, we included behavioural and aging-related modifiers as covariates, such as HIV, substance use, and smoking. Analyses were based on M-values (logit-transformed β-values) to improve statistical properties.

For DMPs, summary statistics from each individual EWAS, segregated by tissue type, were first adjusted for bias and inflation using the empirical null distribution approach implemented in the *bacon* package. The adjusted results were then meta-analyzed using an inverse-variance fixed-effects model in METAL, allowing for effect size pooling across studies. Only CpG sites present in at least three datasets were retained in the meta-analysis to ensure reliability across platforms. Age-associated DMPs were identified using a stringent false discovery rate (FDR) threshold of <0.005.

To examine the directionality of age-related change (hypo-vs. hypermethylation), we combined all datasets available for each tissue and defined methylation categories based on average methylation in young (<30) and old (>60) individuals: High methylation: β ≥ 0.75, Low methylation: β ≤ 0.25, Intermediate: β ≥ 0.25 & β ≤ 0.75 (remaining CpGs). This allowed classification of DMPs according to methylation drift trajectories over time (e.g., high-to-intermediate, intermediate-to-low).

### Variance Analysis (VMPs)

To detect CpGs where the variance of DNA methylation changes systematically with age (VMPs), we employed a two-step Breusch–Pagan heteroscedasticity testing framework. This test evaluates whether the residual variance of DNA methylation is dependent on age, beyond what is explained by standard covariates.

**Step 1:** Fit Linear Models to Estimate Residuals (DMPs analysis)

For each dataset, we first fitted a multivariate linear model for each CpG site, where the dependent variable was the M-value (logit-transformed β-value) of methylation at that CpG. The model was specified as:

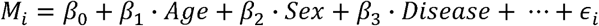

- *M_i_* =*M*-value at CpG site i
- Covariates included:

◦ Age (main variable of interest)
◦ Sex, BMI, technical batch, and other dataset-specific covariates if available
◦ Tissue type was not included as all analyses were stratified by tissue, and meta-analyses were performed within tissue type
- When datasets involved repeated measures (e.g., twins, longitudinal data), a random effect was incorporated using *duplicateCorrelation* from the *limma* package^60^.

This regression model adjusts for known confounders and produces residuals—the component of variation in methylation not explained by age or other covariates.

**Step 2:** Breusch–Pagan Test to Assess Variance Change with Age

Next, for each CpG site, we squared the residuals from Step 1 and used these as the dependent variable in a second univariate regression, where age was the sole predictor:

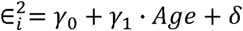

This models whether the variance of methylation residuals increases or decreases with age, i.e., whether age explains variability in methylation, not just average level. CpGs where squared residuals were not normally distributed (Shapiro–Wilk test, p < 0.05) were excluded to avoid inflated Type I error due to residual SNP effects or technical noise.

For VMPs, the output test statistics (χ² values) from the Breusch–Pagan tests were meta-analyzed across datasets using a sample-size-weighted fixed-effects approach, also implemented in METAL. This method accounted for the differing sample sizes among datasets without requiring effect size estimates. CpGs were included in the meta-analysis if they were present in at least 15% of all available samples, and significant age-associated VMPs were defined at an FDR <0.005.

### Entropy Analysis

To quantify methylation stochasticity and epigenetic complexity, Shannon entropy was calculated per sample across multiple sets of CpGs:

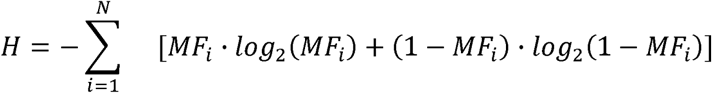

- MF_i_: methylation fraction (β-value) of CpG i
- Entropy ranges from 0 (fully methylated or unmethylated) to 1 (maximal uncertainty at ∼50% methylation)

To isolate age-related effects, we conducted a multivariate linear model regressed M-values against available covariates for each dataset (i.e. sex, BMI, disease, etc. – excluding age). We then added residuals to the mean signal to reconstruct adjusted β-values. Entropy was then computed on these adjusted β matrices, preserving age effects while removing confounders. Age was then regressed against sample-level entropy in each dataset to assess age-dependency:

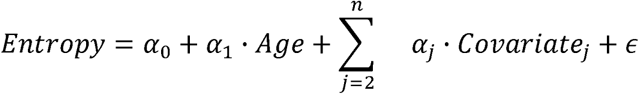

Entropy was computed in a genome-wide manner, across DMPs, VMPs, non-age-associated CpGs.

For entropy, regression coefficients and standard errors from the linear models of age versus genome-wide methylation entropy (and other entropy subsets) were extracted for each dataset and meta-analyzed using a fixed-effects model implemented in the metafor package in R. Separate meta-analyses were performed for entropy calculated over all CpGs (genome-wide), over specific subsets including DMP-only, VMP-only, the intersection of DMPs and VMPs (DMP∩VMP), and CpGs not associated with age.

### Cross-Tissue Integration of DMPs and VMPs

To evaluate the extent of cross-tissue conservation of age-associated epigenetic signatures, we aggregated significant DMPs and VMPs across tissues into binary presence–absence matrices. Jaccard similarity indices and effect size correlation matrices were computed to assess overlap and directional consistency of DMPs and VMPs across tissues. Shared CpGs were defined as those present in multiple tissues, and unique CpGs were analyzed for tissue-specific relevance. A combined similarity heatmap, incorporating Jaccard index (bottom triangle) and effect direction consistency (top triangle), was used to visualize tissue relationships. Permutation testing (10,000 iterations) was employed to determine whether observed DMP overlaps exceeded random expectations, and significance was denoted using empirical p-values. Shared and unique DMP sets were subjected to GO enrichment analysis using missMethyl^61^. Additional integrative visualizations included effect size matrix clustering and annotation of total and overlapping DMPs per tissue. These results enabled robust comparison of conservation, specificity, and convergence of aging-related epigenetic signatures across tissues

### Sensitivity Analyses

To assess the influence of age range and sample characteristics on DMP and VMP discovery, we computed pearson correlations between the number of significant hits and age distribution metrics (mean, median, range) per tissue. Tissues with broader age ranges exhibited higher DMP/VMP detection power, underscoring the importance of age variability in epigenetic studies. Scatterplots with regression and correlation statistics were generated to visualize these relationships. This sensitivity analysis confirmed that observed differences in DMP counts are partially attributable to differences in study design rather than purely biological variation.

### Weighted Gene Co-methylation Network Analysis (WGCNA)

We implemented WGCNA on each tissue individually restricted to age-associated CpGs to identify modules of co-methylated CpGs. Prior to network construction, datasets were harmonized by aligning sample IDs between beta matrices and phenotype metadata, filtering out samples and CpGs with excessive missing values (>10% and >25%, respectively), and removing low-variance CpGs (lowest 20%). We followed the standard framework of WGCNA^62,63^. Initially, we determined the ideal soft-thresholding power for each tissue to uphold a scale-free topology in the methylation co-expression network The soft-thresholding power was selected based on scale-free topology (R² ≥ 0.8) where possible and reasonable mean connectivity. Modules were identified using hierarchical clustering of topological overlap matrices with dynamic tree cutting, and their eigengenes were correlated with chronological age. Statistically significant modules (p < 0.05) were further analyzed. Annotated CpGs within these modules were mapped to genes using the EPIC array annotation. Gene ontology (GO:BP) enrichment was performed for each significant module using enrichGO (clusterProfiler). Modules were then classified based on the direction of association with age (positive, negative), and their top hub CpGs and driver genes were extracted. Enriched GO terms and module–gene interactions were visualized using dot plots and network plots. Shared drivers and their genomic contexts (e.g., promoter, enhancer, gene body) were annotated, and chromatin state distributions were examined using Roadmap Epigenomics data.

### Gene-level Functional Meta-analysis (mitch)

DMP-level t-statistics were aggregated to the gene level using a sum-of-t statistics approach with multiple gene mappings per CpG parsed and split. The resulting gene × tissue t-statistic matrix was imported into the mitch framework for multivariate enrichment testing. Gene sets were sourced from MSigDB (GO:BP), and tested for coordinated enrichment using MANOVA-based statistical models. Pathways significantly enriched (FDR < 0.005) across tissues were classified by effect direction (e.g., s.dist). Both significance-prioritized and effect-size-prioritized runs were performed. Summary statistics and visualizations (bar plots, scatterplots, heatmaps) were generated for downstream interpretation.

### Pan-Tissue Gene-Gene Correlation Network Analysis

The mitch-derived t-statistic matrix was z-score transformed, filtered for genes with low missingness, and converted into a pairwise Spearman correlation matrix. Hierarchical clustering (average linkage) was used to generate gene dendrograms, followed by static (k = 10 and k = 30) and adaptive dynamic tree cutting (deepSplit = 2, minClusterSize = 50) to define gene modules. Directional consistency across tissues was determined for each module (e.g., universal gain/loss, tissue-specific, divergent), and average t-statistics per module-tissue combination were visualized via heatmaps. Top hub genes (highest connectivity) and top driver genes (highest combined connectivity × average t-statistic) were identified for each module. Gene set enrichment for each module was performed using clusterProfiler with MSigDB terms.

### In Silico Robustness Simulations

We developed an in-silico perturbation framework to quantify the impact of methylation shifts in individual genes on global module structure. Perturbations were simulated by zeroing out (knockdown) or doubling (overexpression) t-statistic values across tissues. For each gene perturbation, average module activity was recalculated and compared to baseline. Disruption was quantified as the summed absolute difference in module activity across tissues. The top genes inducing high disruption were labeled as global disruptors. Simulations were also run for combinations of genes (e.g., PCDHG family). Each module’s disruption profile was annotated for pathway enrichment and labeled as “beneficial” or “harmful” based on whether the disrupted module was enriched in age- or disease-associated pathways. Genes with consistent multi-module influence were flagged as cross-tissue disruptors.

## Acknowledgements

Supported by National Institute on Aging and Hevolution grants to VNG.

## Competing Interests

The Regents of the University of California are the sole owner of patents and patent applications directed at epigenetic biomarkers for which Steve Horvath is a named inventor; SH is a founder and paid consultant of the non-profit Epigenetic Clock Development Foundation that licenses these patents. SH is a Principal Investigator at the Altos Labs, Cambridge Institute of Science, a biomedical company that works on rejuvenation. The other authors declare no conflict of interests.

---

**Supplementary Figure 1.**
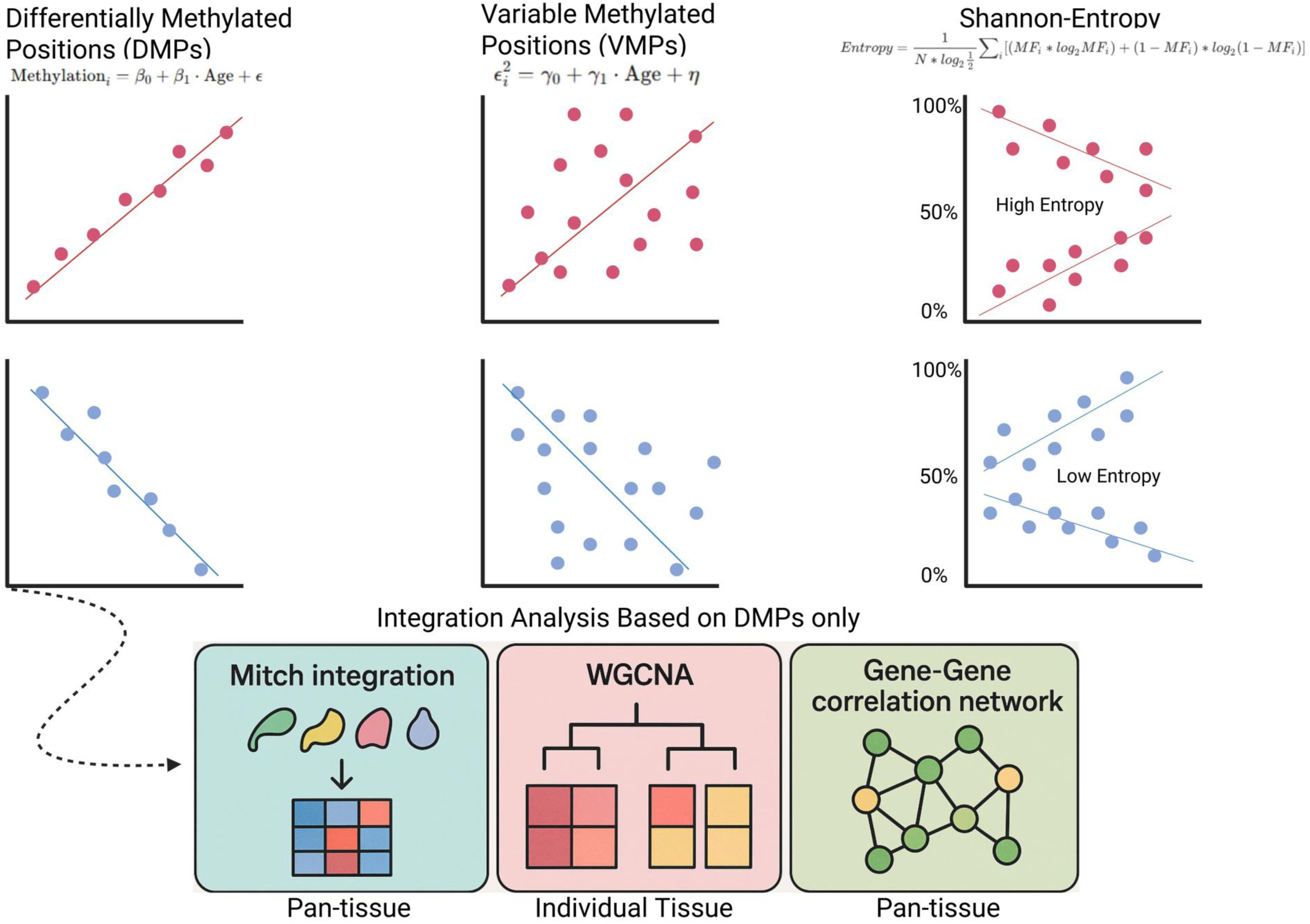
Sensitivity analyses testing **A**: the relationship between VMPs count and cohort characteristics including: sample size (R =, p =); age range (R =, p =); median age (R =, p =); and mean age (R =, p =). **B**: The relationship between DMPs count and cohort characteristics, including: age range (R = 0.66, p = 0.0072); median age (R = 0.22, p = 0.43); and mean age (R = 0.13, p = 0.65). Only the age range was significantly associated with DMP yield, suggesting increased statistical power with broader sampling rather than age-related bias.

**Supplementary Figure 2.**
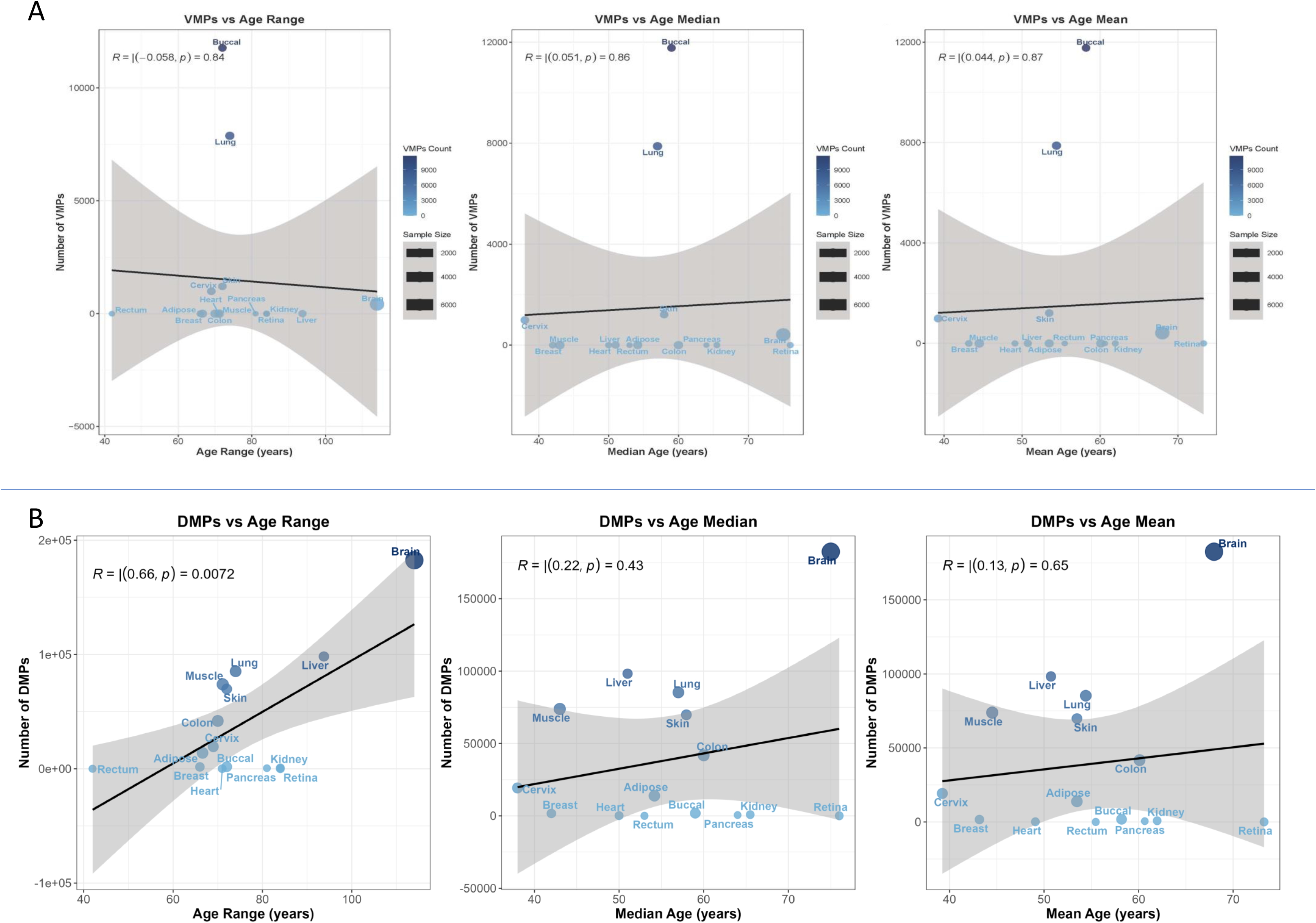
Cross-tissue overlap of age-associated VMPs reveals both conserved and tissue-specific aging signatures. Bottom diagonal half of heatmap showing pairwise Jaccard indices of DMP overlap across human tissues profiled in the aging methylation atlas. Darker blue colors indicate greater overlap, with clustering revealing groups of tissues with shared age-associated DMPs. Most tissues show low overlap, reflecting divergent aging trajectories. Top diagonal half of the heatmap shows effect-size correlation across tissues to determine directionality changes. Darker red colors indicate greater concordance in direction of changes of overlapped VMPs.

**Supplementary Figure 3.**
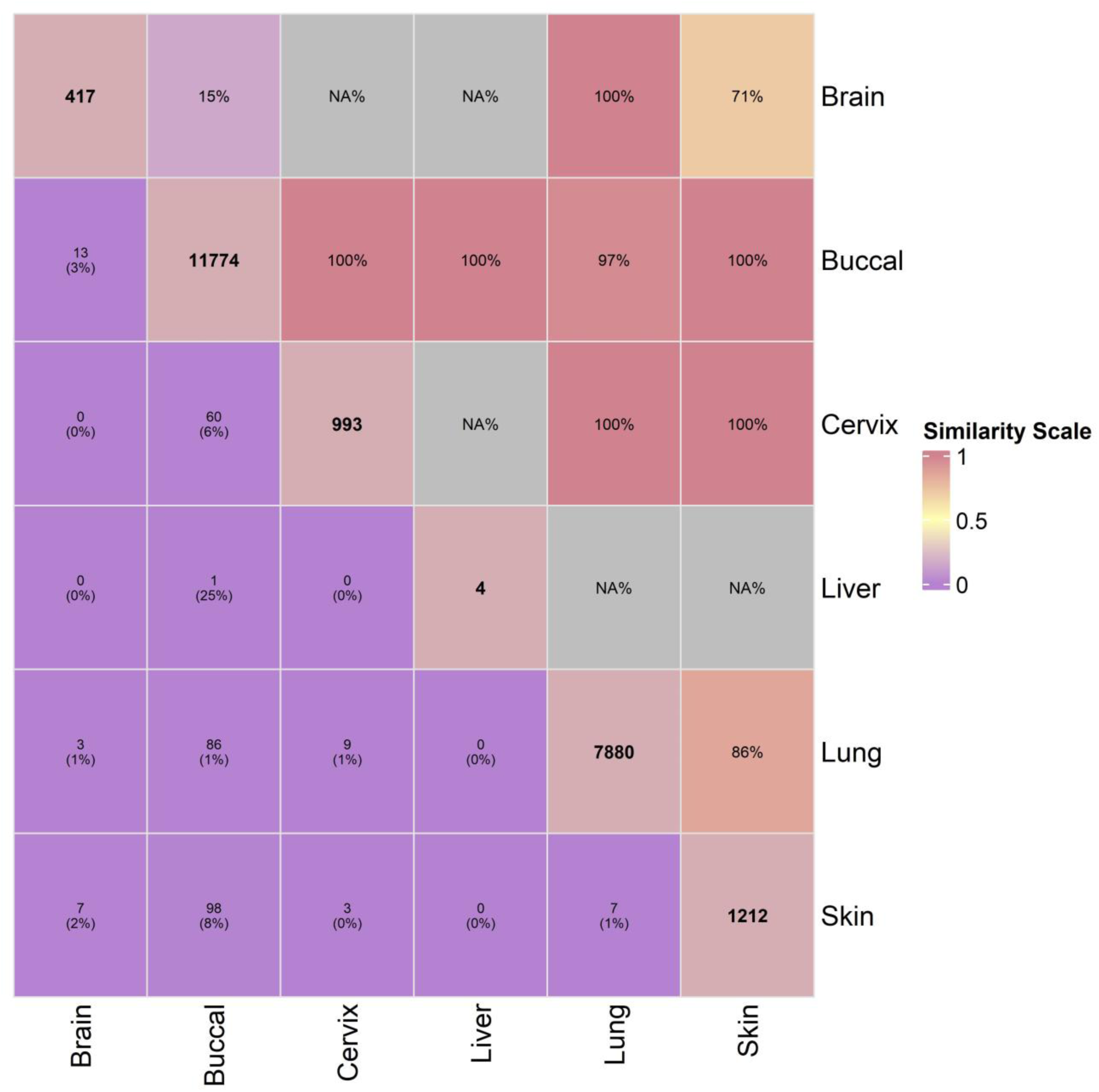
Selection of soft-thresholding power and identification of co-methylation modules using WGCNA. A/C/E/G Scale-free topology fit index (y-axis) plotted against a range of soft-thresholding powers (x-axis) used to determine the optimal power value for network construction. A power value was selected where the scale-free topology fit index approached or exceeded 0.85, indicating a robust scale-free network structure. Mean connectivity (y-axis) plotted against the same range of soft-thresholding powers, illustrating the trade-off between network sparsity and connectivity. B/D/F/H Hierarchical clustering dendrogram of CpGs grouped by topological overlap, with module colors representing distinct co-methylation modules identified by WGCNA. Each branch corresponds to a module of highly correlated CpGs.

**Supplementary Figure 4.**
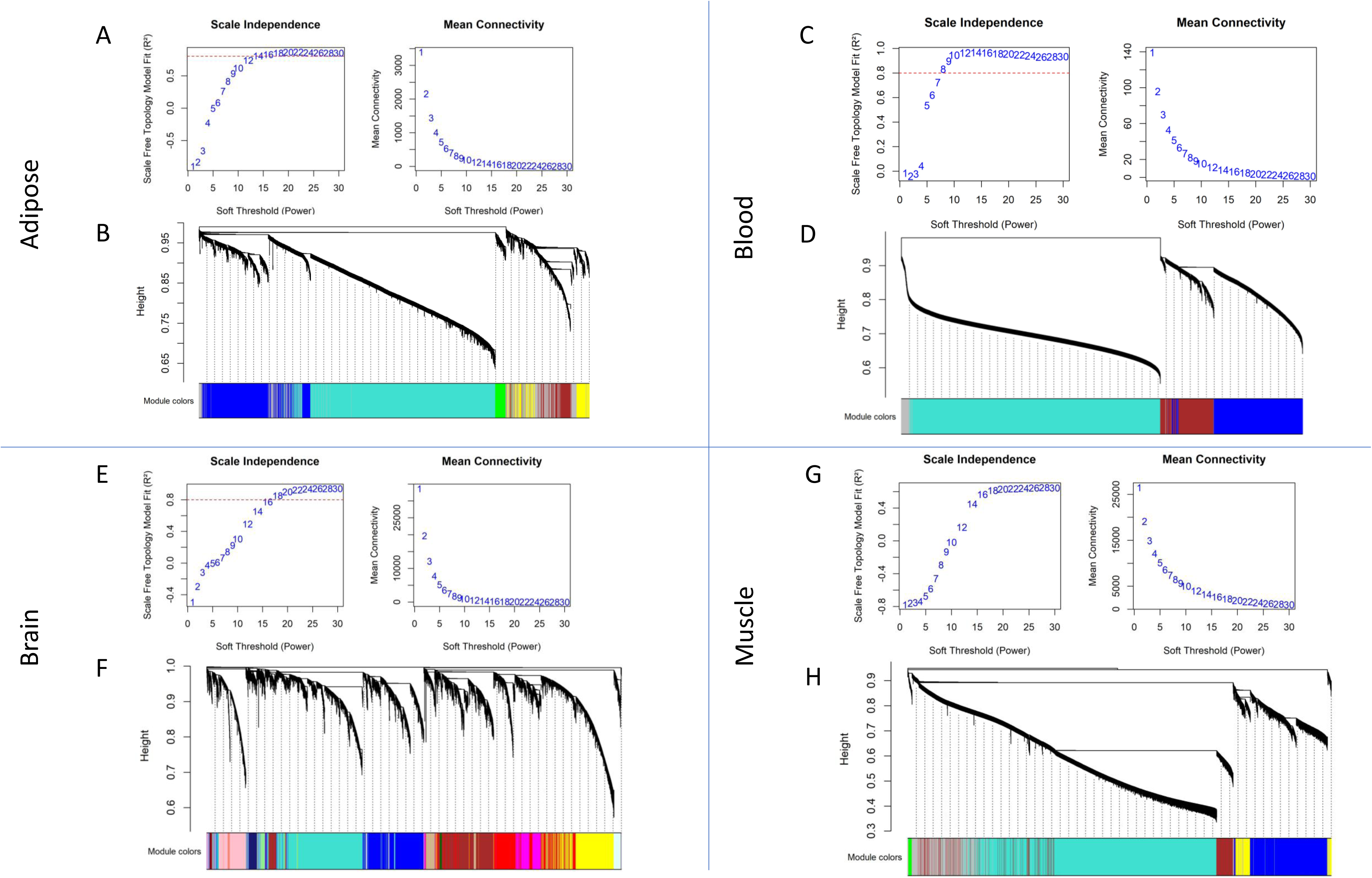
Module directionality and enrichment A/C/E/G Module Correlation for each tissue with directionality. B/D/F/H Top enriched pathways in each module, separated by negative and positive modules.

**Supplementary Figure 5.**
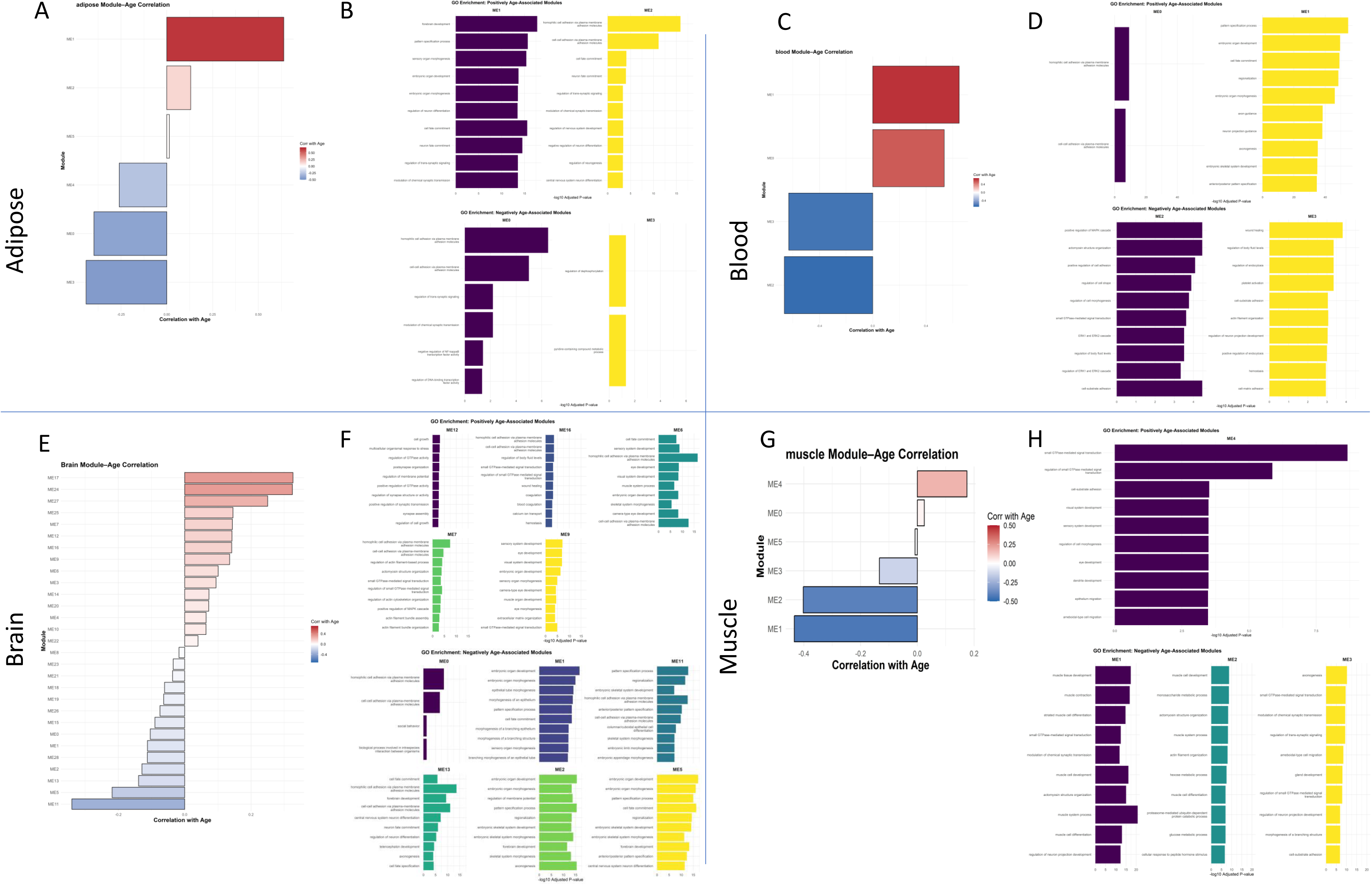
Cross-tissue perturbation analysis network of universal module (Module 5). Driver genes are shown in red (high connectivity × strong t-statistic signal – likely to have functional impact), while hub genes (correlation strength to other genes in the module-seen as coordinators or integrators) are shown in blue.

**Supplementary Figure 6.**
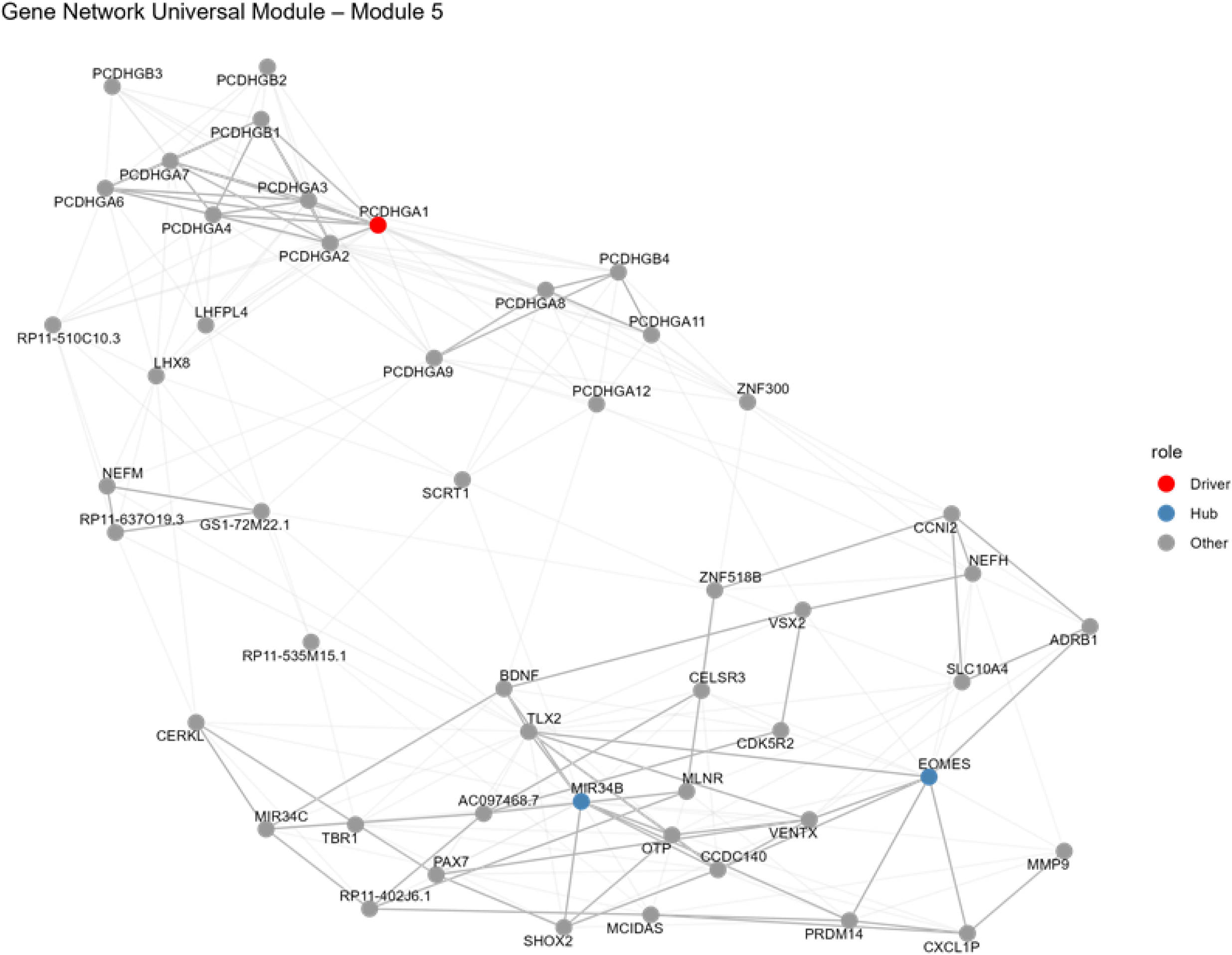
Cross-tissue intersection and Perturbation Analysis

**Supplementary Table 1** Permutation-based sensitivity analysis for the overlap likelihood of VMPs and DMPs.

**Supplementary Table 2** Pan-tissue analysis of gene-gene networks

**Supplementary Table 3** In-silico perturbation analysis findings featuring key contributing genes and pathways

## Notes

https://eynon-lab.shinyapps.io/humanagingatlas

